# gencore: An Efficient Tool to Generate Consensus Reads for Error Suppressing and Duplicate Removing of NGS data

**DOI:** 10.1101/501502

**Authors:** Shifu Chen, Yanqing Zhou, Yaru Chen, Tanxiao Huang, Wenting Liao, Yun Xu, Zhicheng Li, Jia Gu

## Abstract

**Background:** Removing duplicates might be considered as a well-resolved problem in next-generation sequencing (NGS) data processing domain. However, as NGS technology gains more recognition in clinical applications (i.e. cancer testing), researchers start to pay more attention to its sequencing errors, and prefer to remove these errors while performing deduplication operations. Recently, a new technology called unique molecular identifier (UMI) has been developed to better identify sequencing reads derived from different DNA fragments. Most existing duplicate removing tools cannot handle the UMI-integrated data. Some modern tools can work with UMIs, but are usually slow and use too much memory, making them not suitable for cloud-based deployment. Furthermore, existing tools rarely report rich statistical results, which are very important for quality control and downstream analysis. These unmet requirements drove us to develop an ultra-fast, simple, little-weighted but powerful tool for duplicate removing and sequence error suppressing, with features of handling UMIs and reporting informative results.

**Results:** This paper presents an efficient tool *gencore* for duplicate removing and sequence error suppressing of NGS data. This tool clusters the mapped sequencing reads and merges reads in each cluster to generate one single consensus read. While the consensus read is generated, the random errors introduced by library construction and sequencing can be removed. This error-suppressing feature makes *gencore* very suitable for the application of detecting ultra-low frequency mutations from deep sequencing data. When unique molecular identifier (UMI) technology is applied, *gencore* can use them to identify the reads derived from same original DNA fragment. *gencore* reports statistical results in both HTML and JSON formats. The HTML format report contains many interactive figures plotting statistical coverage and duplication information. The JSON format report contains all the statistical results, and is interpretable for downstream programs.

**Conclusions:** Comparing to the conventional tools like Picard and SAMtools, *gencore* greatly reduces the output data’s mapping mismatches, which are mostly caused by errors. Comparing to some new tools like UMI-Reducer and UMI-tools, *gencore* runs much faster, uses less memory, generates better consensus reads and provides simpler interfaces. To our best knowledge, *gencore* is the only duplicate removing tool that generates both informative HTML and JSON reports. This tool is available at: https://github.com/OpenGene/gencore

## Introduction

High-depth next-generation sequencing (NGS) has been widely used for precision cancer diagnosis and treatment [1]. From such deep sequencing data, somatic mutations can be detected to guide personalized targeted therapy or immunotherapy. Recently, circulating tumor DNA (ctDNA) sequencing has been recognized as a promising biomarker for cancer treatment and monitoring. Since the tumor-derived DNA is usually a small part of the total blood cell-free DNA, the mutant allele frequency (MAF) of a variant detected from ctDNA sequencing data can be very low (as low as 0.1%). To detect such low-frequency variants, we usually increase the sequencing depth (can be higher than 10,000x). However, the processes of making NGS library and sequencing are not error-free. Particularly, the library amplification using PCR technology can lead to particular sequences becoming overrepresented [2], and consequently cause some false positive mutations in the result of NGS data analysis.

As a result of library amplification, NGS data can have many duplicates. Especially for the high-depth data generated by sequencing low-input DNA, the duplication level can be much higher. Traditionally, we just mark the duplicated reads and remove them before downstream analysis. For low-depth paired-end NGS data, the read pairs of same start and end mapping positions can be treated as duplicated reads derived from a same original DNA fragment [3]. Then, the reads clustered together can be merged to be a single read. Due to the nature that errors usually happen randomly, the inconsistent mismatches in the clustered read group can be removed to generate a consensus read.

However, for ultra-deep sequencing, it’s possible that two read pairs with same positions are derived from different original DNA fragments. This possibility can be higher when the DNA fragments are shorter. For example, cell-free DNA usually has a peak length of ∼167 bp, which is much shorter than the peak length of normally fragmented genomic DNA. To better identify sequencing reads derived from different DNA fragments, a technology called unique molecular identifier (UMI) has been developed. It has been adopted by various sequencing methods such as Duplex-Seq [4] and iDES [5]. With UMI technology, each DNA fragment is ligated with unique random barcodes before any DNA amplification process. The UMIs can be then used for accurate clustering of sequencing reads. UMIs may be applied to almost any sequencing method where confident identification of PCR duplicates by alignment coordinates alone is not possible and/or an accurate quantification is required, including DNA-seq karyotyping [6] and antibody repertoire sequencing [7].

Some tools, like SAMtools [8] and Sambamba [9], are commonly used to remove duplicates, but cannot process data with UMIs. Samtools is not efficient since it has to sort the data twice for marking duplicate alignments. Sambamba runs faster, but opens a lot of files (much more than 1024), and may introduce problems when multiple instances are run concurrently. The conventional tool Picard markDuplicates (http://broadinstitute.github.io/picard) is able to handle UMIs, but cannot process bam data with unmapped reads. UMI-Reducer [10] and UMI-tools [11] are two new tools designed for processing UMI-integrated NGS data. However, UMI-Reducer is only suitable for RNA data, and UMI-tools cannot deal with data without UMI-integrated. Furthermore, these tools are usually relative slow and use too much memory, which make them cost ineffective for cloud-based deployment. These unmet requirements drove us to develop a new tool called *gencore*, which is fast and memory efficient, with functions to eliminate errors and remove duplicates by generating consensus reads for NGS data with or without UMIs. Table 1 shows a brief comparison of features of different deduplication or consensus read generating tools.

**Table 1.** Features comparison of different deduplication or consensus read generating tools.

|  | SAMtools | Picard | *gencore* | Picard | UMI-tools | *gencore* |
| --- | --- | --- | --- | --- | --- | --- |
|  | Non-UMI mode |  |  | UMI mode |  |  |
| No need to sort by read name |  | + | + |  | + | + |
| No need to sort clean up flags |  | + | + |  | + | + |
| No need to sort add UMI tag | + | + | + |  | + | + |
| No need to sort by position again |  | + | + |  | + | + |
| JSON Report | + | + | + | + |  | + |
| HTML Report |  |  |  |  |  | + |

In Table 1, the input for these five tools is a sorted bam. SAMtools cannot handle UMIs, whereas UMI-tools is only applicable for UMI-integrated data. Only UMI-tools and *gencore* needn’t any extra BAM preporcessing before performing the deduplication. *gencore* reports metrix in JSON and HTML formats whereas UMI-tools doesn’t.

## Implementation

*gencore* requires an input of position sorted BAM file and a reference genome FASTA file. If the FASTQ data has UMIs, it can be preprocessed using fastp [12] to move the UMIs from read sequences to read identifiers. The main workflow of *gencore* is described in Fig.1.

**Fig.1.**
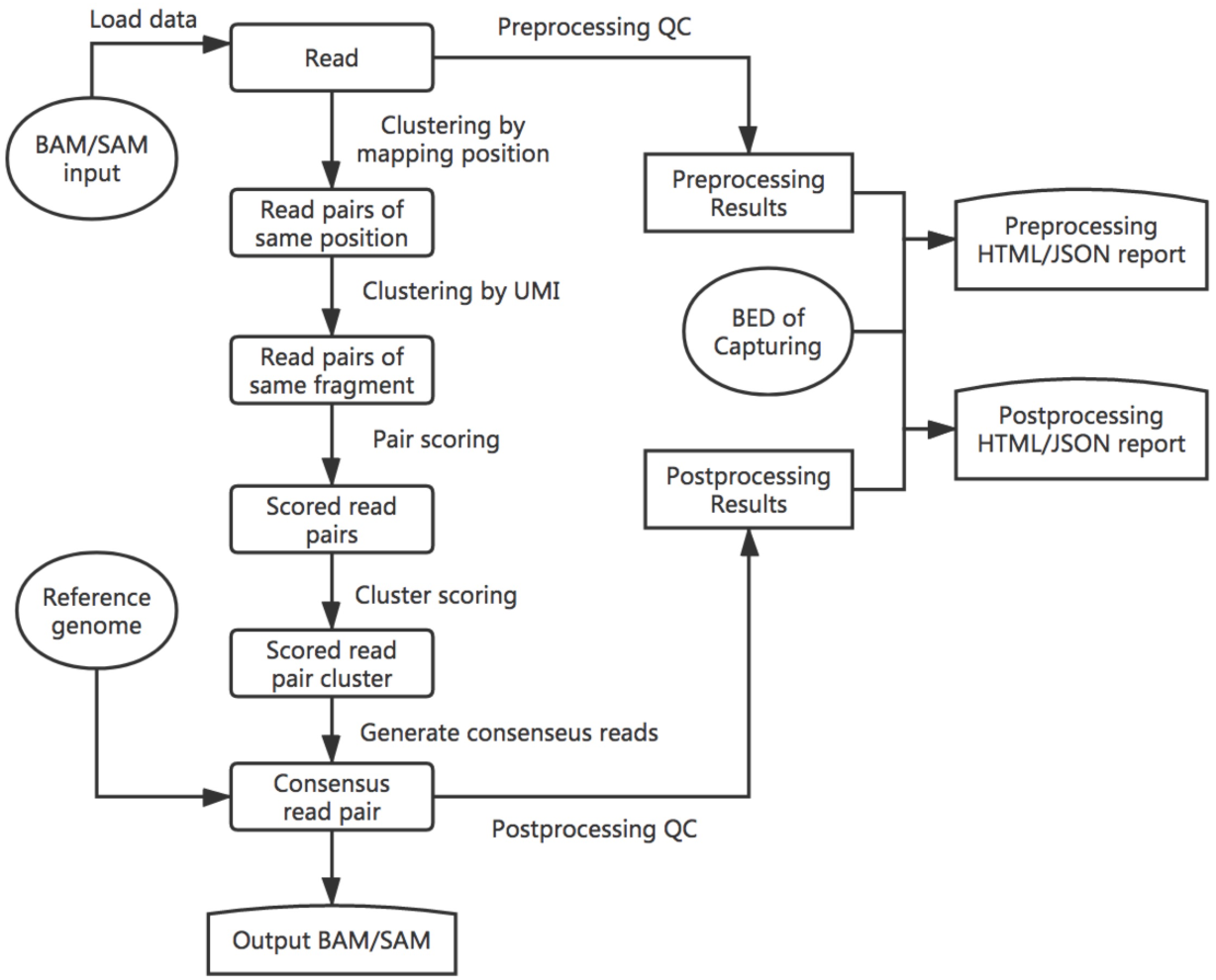
The brief workflow of *gencore*. Besides the input BAM/SAM file, this tool accepts a reference genome input to assist consensus reads generation. If the data is from targeted sequencing, a BED file can also be provided to describe the capturing regions. In this case, the coverage statistics in BED regions will also be reported in the HTML/JSON reports.

Simply put, *gencore* clusters read pairs by their mapping positions and UMIs (if applicable), and then generates a consensus read for each cluster. The main algorithm of *gencore* can be briefly introduced as following steps:

1. Position clustering: all mapped read pairs are grouped by mapping position first. The reads with same mapping chromosome, start position and end position will be grouped together. A multi-level map [*chr*]:[*left_pos*]:[*right_pos*]:[*read_pairs*], is used to store the clusters being processed, while [*left_pos*] and is [*right_pos*] the read pair’s leftmost and rightmost mapping position in the chromosome respectively. [*read_pairs*] is a group of read pairs that share the same mapping positions. To reduce the memory usage, *gencore* implements a processing-while-reading strategy, which means processing one group immediately when its all possible reads are collected. For example, when *gencore* finds that the mapping position of current inputting read is greater than [*right_pos*] of one group, it will perform following step 2 to step 7 for this group and release the group immediately.
2. UMI clustering: for each group of same mapping positions, read pairs are then clustered by their UMIs with one base difference tolerance. If the data has no UMIs, this step is skipped. Due to the principle of Illumina paired-end sequencing, if the data has dual UMIs from forward and reverse reads, the read pairs with reciprocal UMIs will be clustered together. For instance, two read pairs with UMI *ATGC_GCAA* and *GCAA_ATGC* will be considered as derived from different strands of same original DNA fragment, and will be clustered together.
3. Cluster filtering: each cluster will be filtered by comparing its supporting reads number with the threshold (default = 1, which means no threshold). If it passes, *gencore* will start to generate a consensus read for this cluster. For ultra-deep sequencing (i.e. ctDNA sequencing with 10,000× or higher depth), it’s recommended to increase the threshold to 2 to discard part of reads that without any PCR duplicates.
4. Pair scoring: a default score number (default = 6) will be initially assigned to every base in the read. For each read pair in a cluster, the overlapped region of the paired reads is computed. For each base in the overlapped region, its score is adjusted according to its consistence with its paired base, with the consideration of their quality scores. The default scoring schema is presented in the project repository, and can be configured through options.
5. Cluster scoring: in this step, the total scores are computed by summarizing the scores computed in previous step. For each position in the mapping region, *gencore* queries the base presented in the cluster’s different reads, and summarizes them to compute the score of different bases (A/T/C/G).
6. Consensus read generating: for each position in a cluster, its base diversity is computed according to the scores of different bases computed in last step. If *gencore* finds one dominant base, this base will also be presented in the consensus read. Otherwise if exists one or more reads are concordant with reference genome with high quality, or all reads at this positions are with low quality, the corresponding base in the reference genome will be used. The using of reference genome is one of the major differences between *gencore* and other tools. Since a base is more likely an error when it’s not concordant with reference, *gencore* assigns lower weight to them when computing the consensus reads.
7. Buffered reads outputting: when one consensus read is generated, it will be buffered in a position-sorted queue to be written to output. To minimize the memory used by this queue, *gencore* implements a writing-while-processing strategy. With this strategy, *gencore* maintains a pointer that always points to the unprocessed read with least mapping position, and periodically outputs the reads in queue with mapping position less than it.

After the processing is done, *gencore* will generate a summary of the data before and after processing. Some metrics like coverage, duplication histogram, mapping rate, duplication rate, passing filter rate and mismatch rate are reported in HTML/JSON format reports. The HTML report contains no image figures but some interactive figures, which are built based on Plotly.js. Comparing to conventional HTML reports with static images, this single-page standalone JavaScript-based HTML report is much more interactive and easier to transfer. Fig. 2 shows a demonstration of the coverage statistics in both genome scale and capturing regions in the HTML reports.

**Fig. 2.**
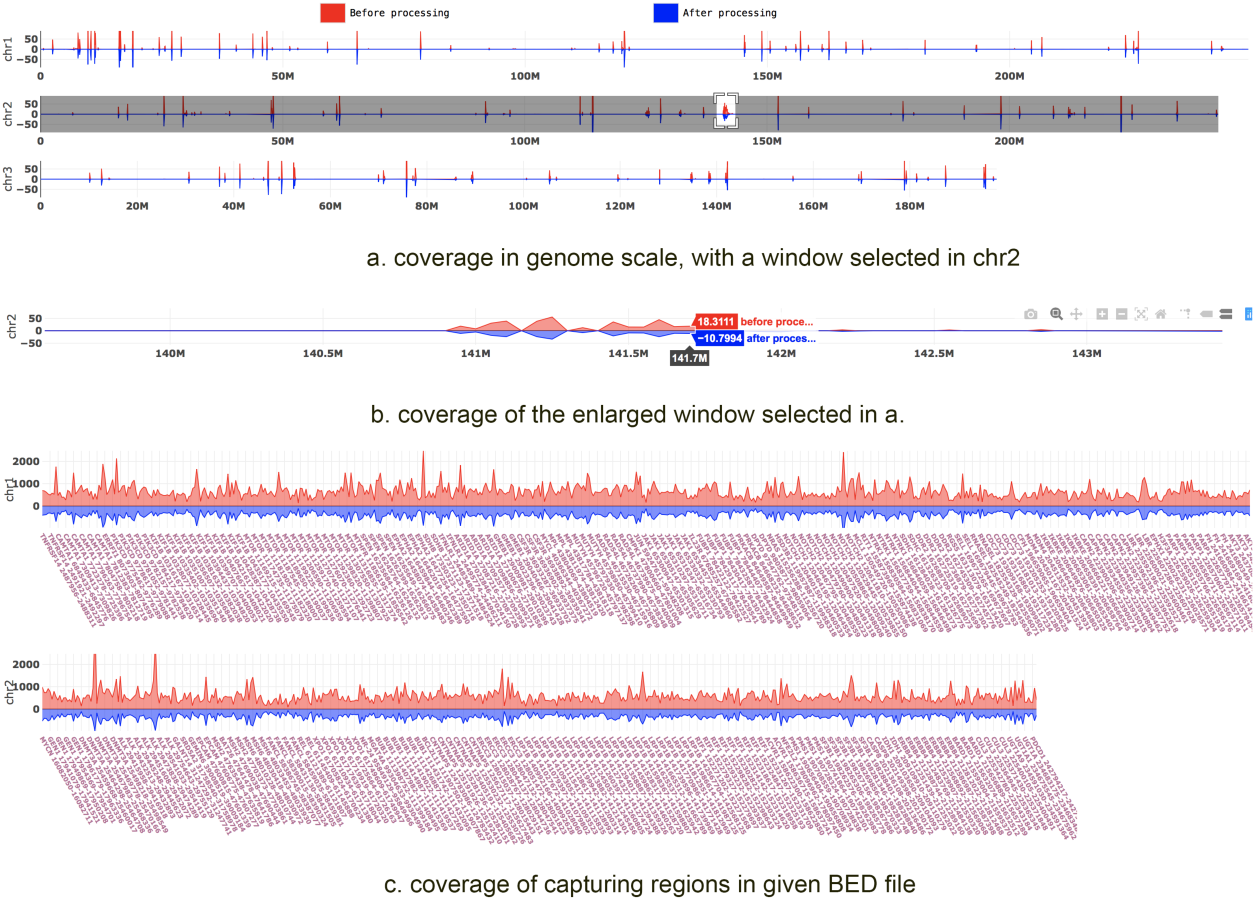
The coverage statistics figures in the HTML report. In this interactive HTML report, a region is selected in a), and then enlarged in b). While a) and b) are coverage in genome scale, c) is the coverage only in the capturing regions. The BAM file of this report was generated by targeted sequencing using a panel with hundreds of genes. So the coverage in genome scale is very sparse, whereas the coverage in capturing regions is high and dense.

## Application

Since *gencore* can be used to reduce sequencing errors, it is very useful for the application of detecting low-frequency somatic mutations from cancer sequencing data, particularly in liquid biopsy technology [16]. When the samples are from blood, urine or malignant effusion, the MAF of variants can be even much lower than 1%. The detection of such low-frequency variants can be seriously affected by the errors, which are usually introduced by library preparation and sequencing. *gencore* can significantly reduce the sequencing errors of deep sequencing data, and consequently reduce the false positive calling rate.

To evaluate how *gencore* can help the variant detection, we conducted two evaluation experiments using 8 DNA samples, obtained from the National Center for Clinical Laboratories (NCCL) in China. The dataset #1 (1801, 1802, 1803 and 180N) was generated by sequencing plasma cell-free DNA samples, and each data contains ∼55G bases. Sample 1801, 1802 and 1803 were DNA extracted from blood of one lung adenocarcinoma patient at different time, whereas sample 180N was DNA from a healthy control. The dataset #2 (1811, 1812, 1813 and 181N) was generated by sequencing tissue DNA samples, and each data contains ∼10G bases. Sample 1811 and 1812 were DNA of tumor tissues collected from two breast cancer patients, whereas sample 1813 was DNA of a biopsy tissue collected from a lung adenocarcinoma patient, and sample 181N was DNA from a healthy control. These DNA samples are publicly provided by NCCL as reference materials for conducting inter-lab quality assessments. The golden standard mutations of all samples were also provided by NCCL. Among all the mutations, the lowest frequency was about 0.15%. In our experiment, all samples sequencing libraries were prepared using IDT xGen Dual Index Adapters, captured with a 451-gene cancer panel, and then sequenced using an Illumina NovaSeq 6000 sequencer. UMI adapters were used for 1801, 1802, 1803 and 180N samples. The detailed file sizes and commands of experiments are provided in Supplementary file 1.

The FASTQ files were preprocessed by fastp, and then mapped to reference genome hg19 using BWA [13]. After the mapped bam file was sorted using Samtools, the sorted bam files were then processed by different tools. VarScan2 [14] was used to call SNVs from the processed bam files, and ANNOVAR [15] was then used to annotate the VCF files. The missense variants detected in the coding sequences were then filtered with conditions (dataset #1: supporting reads ⩾ 5; dataset #2: supporting reads ⩾ 8 and variant allele frequency ⩾ 2%). The variant calling results were evaluated by comparing to the golden standard results provided by NCCL, and the speed and memory performance were also compared. The comparison result is shown in Fig 3.

**Fig. 3.**
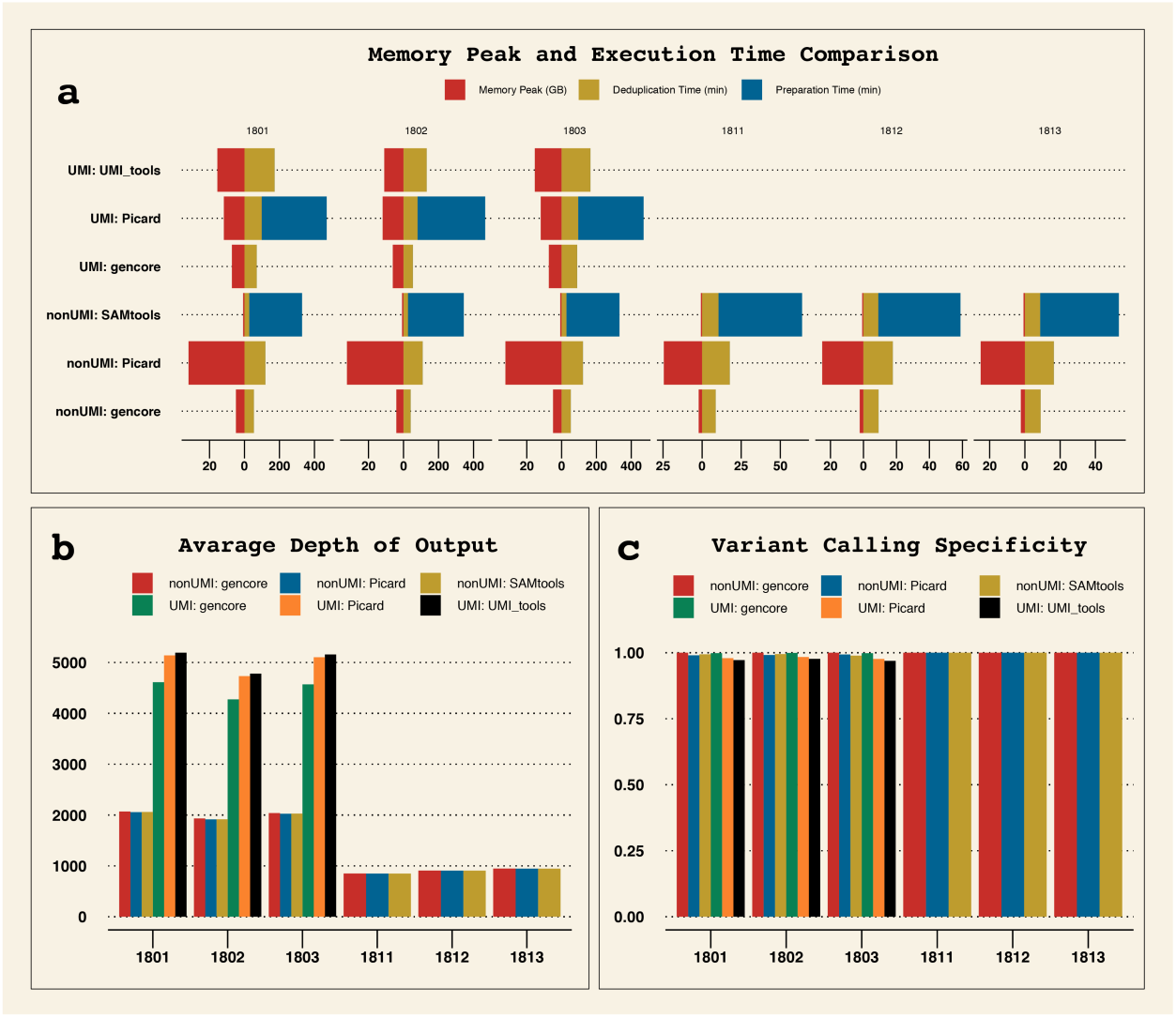
Comparison of speed, memory peak and processing results of different tools in both UMI and non-UMI modes. a) memory peak and execution time of different tools. Samtools and Picard (in UMI mode) need to prepare the data before performing deduplication, whereas *gencore* and UMI_tools needn’t. b) average depth of output BAM. For the cfDNA samples (1801, 1802 and 1803), the depths of UMI mode results are much higher than non-UMI mode, indicating that over-deduplication may happen when performing deduplication without UMI for ultra-deep sequencing data. c) specificity of downstream variant calling results comparing to the golden standard results provided by NCCL.

From Fig. 3, we can learn that *gencore* runs much faster than all other tools. For the comparison of memory peak, *gencore* uses much less memory than Picard and UMI_tools. Due to *gencore* consumes extra memory to load reference genome and performs more processing, *gencore* uses more memory than Samtools. But, as shown on Fig. 3a, its memory peak is still less than 8GB. This result shows that *gencore* is littleweight and fast, and is much more cost-effective to be deployed on the cloud. The comparison of downstream variant calling results also shows that *gencore* outperforms other tools. From Fig. 3c, we can learn that *gencore* achieved higher specificity in both UMI and non-UMI modes.

For the cfDNA samples (1801, 1802 and 1803), we applied a filter with condition (supporting reads⩾5). The results showed all tools successfully detected all true positive variants for 1801 and 1802, but non-UMI mode tools missed one true positive variant for 1803 due to its variant allele frequency was too low (VAF = 0.15%). Moreover, we evaluated the detected variants by comparing their VAFs to the golden results, and considered a variant as unacceptable if its VAF exceeded 2 standard deviations. The results showed all non-UMI mode tools resulted in two variants with unacceptable VAFs. For UMI mode, UMI_tools detected one variant with unacceptable VAF, while all variants detected by Picard and *gencore* were acceptable. For the tissue DNA samples (1811, 1812 and 1813), we applied a filter with condition (supporting reads ⩾ 8 and VAF ⩾ 2%). The results showed that all tools could detect true positive variants at 100% sensitivity. But for sample 1811, both Picard and Samtools reported one false positive variant, while *gencore* achieved 100% specificity for all three samples.

These results suggest that UMI technique is important for detecting variants with ultra-low VAFs, and *gencore* is one of the best tools to process UMI-enabled data due to its superior accuracy and performance.

## Results and discussion

By analysing the output data using downstream tools, *gencore* outperforms other tools in both non-UMI and UMI modes. By carefully exploring the data generated by these different tools, we found the major difference was that *gencore* applied reference genome based correction, whereas Picard and UMI-tools didn’t. Utilization of a reference genome is important for eliminating sequencing noises. When an inconsistent position is found when making a consensus read, the reference base should be taken into account since the base different from the reference may have higher probability to be a sequencing error.

To explore how *gencore* eliminates the sequencing errors, we manually compared some the alignment files before and after *gencore* processing. In the case of sample 1802, NM_005228.3(EGFR): c.2369C>T, p.T790M variant was one true positive variant. Fig. 4 shows the alignment visualization illuminated by Integrated Genome Viewer (IGV) for the files before and after processing. In Fig. 4a, which is the original alignment file generated by mapping by BWA [13], the double-line marked mismatch T base is the true positive variant EGFR p.T790M. However, there are also some other mismatch bases, which are false positive mismatches caused by sequencing errors. In Fig. 4b, which is the alignment file after *gencore* processing, we can find these false positive mismatches are gone, while the true positive variant is kept. This result suggests that *gencore* not only removes duplicates, but also eliminates sequencing errors.

**Fig. 4.**
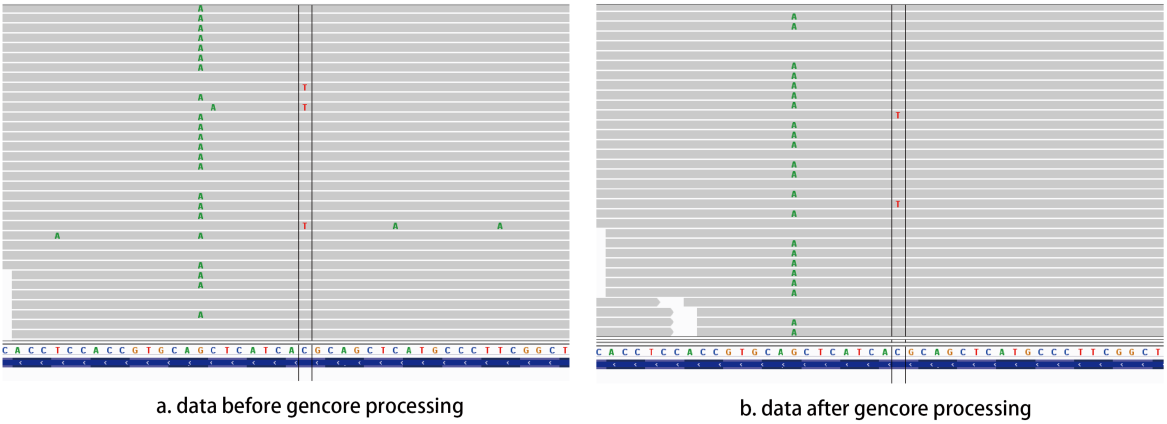
Comparison of the alignment files before and after *gencore* processing. In this figure, the position marked by double lines is NM_005228.3(EGFR): c.2369C>T, p.T790M variant. a) visualizes the mapped reads of original alignment file, b) visualizes the mapped reads after *gencore* processing. We can find that the false positive mismatches, which appear randomly in the original alignment file, are corrected by *gencore*.

## Conclusion

We introduced a tool *gencore*, which is useful for performing deduplication and consensus read generation for deep next-generation sequencing data. We conducted several experiments to evaluate the performance of *gencore*, with comparisons to Picard, Samtools and UMI-tools. The result shows that *gencore* is much faster and more memory efficient, while providing similar or better results. This tool generates interactive HTML reports and informative JSON reports that can help manually checking and programmatically downstream analysis. According to our estimation, this tool has been used to process more than 10,000 samples in the authors’ institution, and is now suitable to be adopted by community users.

## List of abbreviations

ctDNA: cell-free tumor DNA;
NGS: next generation sequencing;
IGV: integrative genome viewer;
MAF: mutated allele frequency;
INDEL: insertion and deletion;
SNP: single-nucleotide polymorphism;
SNV: single-nucleotide variation;
EGFR: epidermal growth factor receptor;
UMI: unique molecular identifier;
HTML: Hypertext Markup Language;
JSON: JavaScript Object Notation;
NCCL: National Center for Clinical Laboratories;

## Declarations

### Ethics approval and consent to participate

Not applicable.

### Consent to publish

Not applicable.

### Availability and requirements

Project name: gencore

Project home page: https://github.com/OpenGene/gencore

Operating system(s): Linux or Mac OS X

Programming language: C++

Other requirements: htslib and zlib

License: MIT License.

### Authors’ contributions

SC designed the algorithm, developed the software and wrote the paper. YZ co-designed the algorithm, YC, YX, WL and TH conducted the testing work and evaluation experiments, ZL and JG contributed to data analysis and paper revision. All authors read and approved the final manuscript.

## Acknowledgments

We’d like to thank HaploX Biotechnology for supporting the publication cost. We’d like to express our thanks to the *gencore* community users for testing and reporting issues. We’d like to thank the OpenGene members for their suggestions of *gencore*’s design.

