## Supplementary file 1 for "gencore: An Efficient Tool to Generate Consensus Reads for Error Suppressing and Duplicate Removing of NGS data"

| Tools | Version | Parameters |
|---|---|---|
| *gencore* | 0.14.0 | UMI mode : default<br>nonUMI mode : -u UMI |
| picard | 2.18.26-SNAPSHOT | nonUMI mode: MarkDuplicates REMOVE_SEQUENCING_DUPLICATES=true<br>UMI mode: UmiAwareMarkDuplicatesWithMateCigar REMOVE_DUPLICATES=true |
| samtools | 1.9 | markdup -r -s |
| umi_tools | 1.0.0 | dedup --paired --umi-separator=: |

| Sample | 1801 | 1802 | 1803 | 180N | 1811 | 1812 | 1813 | 181N |
|---|---|---|---|---|---|---|---|---|
| Fastq data size( Gb ) | 56.7 | 51.8 | 54.8 | 59.2 | 11.1 | 9.9 | 9.6 | 10. |
